# Deep learning uncovers histological patterns of YAP1/TEAD activity related to disease aggressiveness in cancer patients

**DOI:** 10.1101/2024.06.14.598991

**Authors:** Benoit Schmauch, Vincent Cabeli, Omar Darwiche Domingues, Jean-Eudes Le Douget, Alexandra Hardy, Reda Belbahri, Charles Maussion, Alberto Romagnoni, Markus Eckstein, Florian Fuchs, Aurélie Swalduz, Sylvie Lantuejoul, Hugo Crochet, François Ghiringhelli, Valentin Derangere, Caroline Truntzer, Harvey Pass, Andre L. Moreira, Luis Chiriboga, Yuanning Zheng, Michael Ozawa, Brooke E. Howitt, Olivier Gevaert, Nicolas Girard, Elton Rexhepaj, Iris Valtingojer, Laurent Debussche, Emanuele de Rinaldis, Frank Nestle, Emmanuel Spanakis, Valeria R. Fantin, Eric Y. Durand, Marion Classe, Katharina Von Loga, Elodie Pronier, Matteo Cesaroni

## Abstract

Over the last decade, Hippo signaling has emerged as a major tumor-suppressing pathway. Its dysregulation is associated with abnormal expression of *YAP1* and *TEAD*-family genes. Recent works have highlighted the role of YAP1/TEAD activity in several cancers and its potential therapeutic implications. Therefore, identifying patients with a dysregulated Hippo pathway is key to enhancing treatment impact. Although recent studies have derived RNAseq-based signatures, there remains a need for a reproducible and cost-effective method to measure the pathway activation. In recent years, deep learning applied to histology slides have emerged as an effective way to predict molecular information from a data modality available in clinical routine. Here, we trained models to predict YAP1/TEAD activity from H&E-stained histology slides in multiple cancers. The robustness of our approach was assessed in seven independent validation cohorts. Finally, we showed that histological markers of disease aggressiveness were associated with dysfunctional Hippo signaling.

## Introduction

Tumorigenesis and cancer progression usually involve the dysregulation of several signaling pathways, particularly those governing cell proliferation and survival. The Hippo pathway, recognized as a tumor suppressor pathway, is one of the major pathways known to be implicated in tumorigenesis, as it inhibits excess proliferation and regulates organ size and apoptosis evasion^1,2^. Several components of this pathway have been shown to be inhibited in certain types of cancer^3^. The yes-associated protein (YAP1) and the transcriptional coactivator with PDZ-binding motif (TAZ) are two primary effectors of the pathway^4^. The shuffling of the YAP1/TAZ complex between the nucleus and the cytoplasm is controlled by phosphorylation. When phosphorylated by the large tumor suppressor (LATS) family of Hippo kinases^5^, the complex remains in the cytoplasm and is sequestered via its interaction with 14-3-3 proteins^6^ and/or degraded after its ubiquitylation^7^. Once in the nucleus, the YAP1/TAZ complex preferentially binds to the transcriptional enhancer associate domain (TEAD) proteins to increase their transcriptional activity^8^. By activating the transcription of genes involved in cell proliferation and transformation (*CTGF* and *CYR61*, for example), YAP1/TEAD activity promotes tumor growth and resistance to chemotherapy^3^.

In malignant pleural mesothelioma (MESO), a cancer of the pleura specifically associated with asbestos exposure, genes within the Hippo pathway are frequently mutated, suggesting a critical role for this pathway in MESO tumorigenesis^9^. Recently, Calvet *et al.*^10^ have demonstrated that YAP1 downregulation induced tumor growth inhibition in xenograft models of a Hippo deficient-mesothelioma cell line *(MSTO-211H)* expressing dominant-negative *TEAD* genes.

Beyond mesothelioma, aberrant YAP1/TEAD activity is also associated with tumor progression and poor prognosis of lung cancer patients. Notably, the missense mutation R133W of the YAP1 protein has been found to be predisposing for lung adenocarcinoma^11^. LATS1 and LATS2, which are responsible for the inactivation of YAP1 and TAZ through phosphorylation, are downregulated in non-small cell lung cancer (NSCLC) compared to healthy tissues^12^ (Malik et al, 2018). Moreover, abnormalities in the Hippo pathway are also found in various malignancies, including breast cancer^13^, gastric cancer^14^, and hepatocellular cancer^15^.

These recent discoveries have spurred interest in targeting the Hippo pathway as a therapeutic strategy in oncology, with several clinical trials currently underway^16,17^. To ensure these treatments are used in the proper indications and given to the appropriate patient subpopulations, it is crucial to develop methods for measuring pathway activation that are both reproducible, as well as time- and cost-effective in a clinical setting.

Activation of YAP1 and TAZ has previously been used as a biomarker of Hippo pathway inactivation^18^. Recently, a comprehensive molecular characterization of 19 Hippo core genes was performed in 9,125 tumor samples across 33 cancer types using multidimensional “omics” data from The Cancer Genome Atlas (TCGA)^19^. This study characterized genetic regulators of the Hippo pathway and identified downstream target genes associated with Hippo pathway activity. To measure YAP1/TEAD activity in tumors and to evaluate the pharmacodynamic effects of TEAD inhibitors, Calvet and colleagues also defined a transcriptional signature of 500 downstream TEAD-effector genes, the expression of which was modified by various perturbations of the Hippo pathway^10,20^. This signature, referred to as TEAD-500, was effectively modulated by knock-out or knock-down of the transcription factors (*YAP1*, *WWTR1* or *TEAD*), and its score was positively correlated with tumor growth (diminishing with tumor regression) in a human cell line used as xenograft^10^.

However, since histology slides are ubiquitous in clinical practice, the identification of image-based biomarkers could allow for a better and generalizable stratification of patients harboring a high YAP1/TEAD activity. In recent years, machine learning approaches to digital pathology have encountered success in several applications, including diagnosis^21^, prognosis^22^, characterization of the immune tumor environment^23^, prediction of genomic alterations^24,25^ and of gene expression^26^. Some of these tools have already made their way to clinical applications, as exemplified by the FDA approval of the first automatic prostate cancer grading system, Paige Prostate^21^, in 2021.

Here, we developed a deep learning approach to predict the YAP1/TEAD activity, measured by the TEAD-500 signature, from histology slides and assessed its relevance, not only in malignant pleural mesothelioma but in 18 different TCGA indications (Figure 1). The models we trained rely on weakly supervised learning to identify morphological patterns predictive of the signature value. They were then validated on no less than seven external datasets, and histological patterns highlighted by the model were subsequently analyzed by pathologists to better describe histological structures and underlying phenotypes related to TEAD activity.

**Figure 1.**
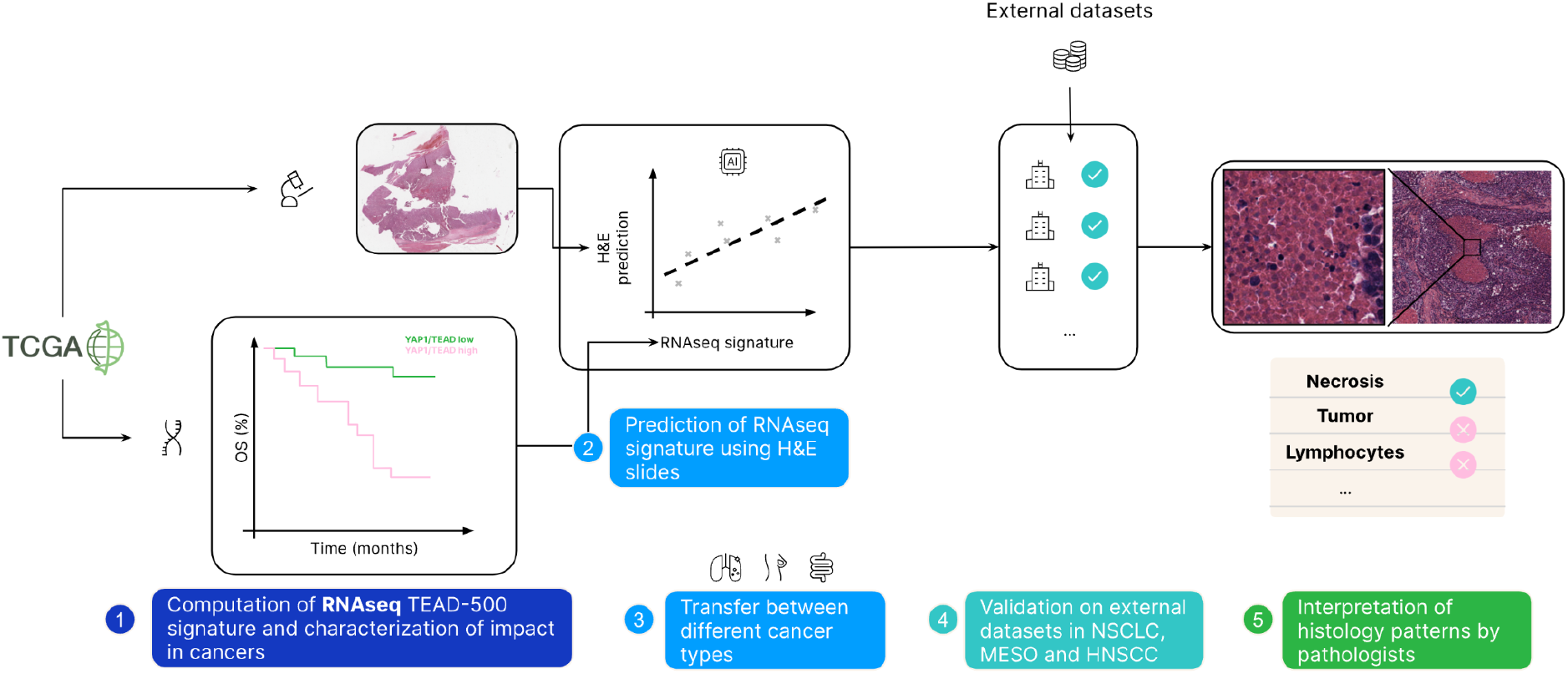
Graphical summary. Graphical abstract. (1) RNAseq and clinical data from 32 cancer types from The Cancer Genome Atlas (TCGA) were collected. TEAD-500 signature was computed from RNAseq data. General statistics and association with prognosis were evaluated. (2) Models were trained to predict the signature from Hematoxylin & eosin (H&E)-stained histology slides. Cross-validation was run on TCGA, and the best models were further validated on seven external cohorts of Non-Small Cell Lung Cancer (NSCLC), mesothelioma and Head and Neck Squamous Cell carcinoma (HNSC), demonstrating the robustness of our approach. (3) The most predictive tiles were extracted from the model and reviewed by trained pathologists to highlight histological biomarkers associated with YAP1/TEAD activity.

## Results

### YAP1/TEAD activity across various cancer indications

We measured YAP1/TEAD activity using a published transcriptional signature of approximately 500 downstream effectors^20^ (referred to herein as the TEAD-500 signature). This signature has been derived from various genetic perturbation experiments and is used as described in Calvet et al^10^ (2022) and summarized in our Methods section. The first goal was to establish a threshold score for YAP1/TEAD activity that could be considered ‘abnormally high’ for any given tumor indication. As expected, the YAP1/TEAD activity and the TEAD-500 score varied widely among different types of normal tissues from The Genotype Tissue Expression project (GTEx) and across TCGA indications (Figure 2a). We therefore defined a distinct threshold per indication, corresponding to the 95^th^ percentile of the score in GTEx samples of the same healthy tissue type. As no comparable healthy tissue collection was available in GTEx for the head and neck squamous cell carcinoma indication (HNSC), non-tumor tissues from the CPTAC-HNSCC cohort^27^ were used as a reference instead. The rationale for choosing the 95th percentile threshold was purely statistical, to match the most common conventional level of significance.

**Figure 2.**
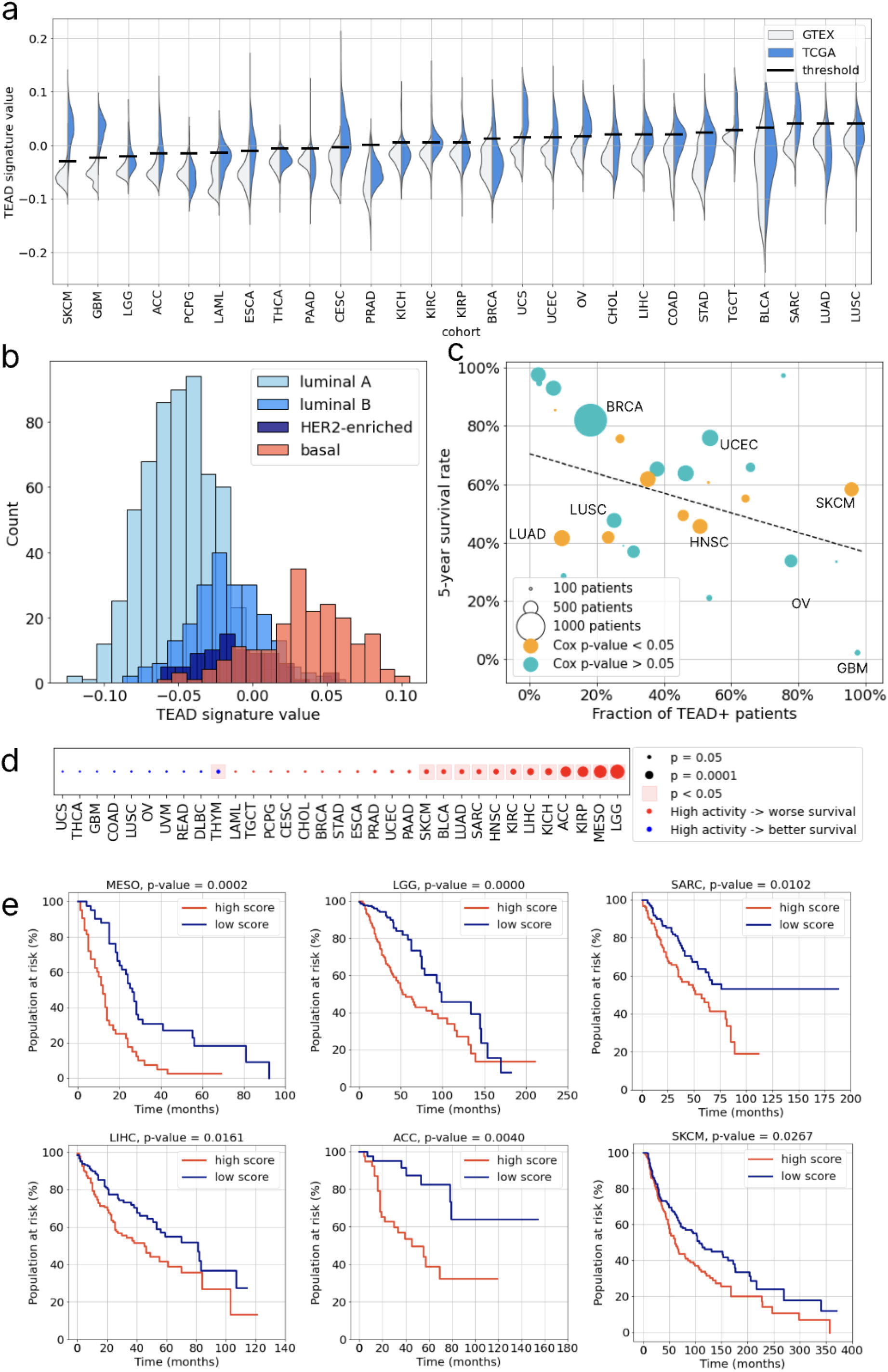
YAP1/TEAD activity is associated with poor prognosis across cancer. **a.** Distribution of TEAD-500 signature values on TCGA cohorts and on matched healthy tissues from the GTEx database. Bars indicate the 95th percentile value on healthy tissue. **b.** Distribution of TEAD-500 signature values in subtypes of BReast CAncer (BRCA). **c.** Correlation between the fraction of patients with a signature value above the GTEx-inferred threshold, and Overall Survival (OS). In orange, cohorts where the Cox p-value of the signature is significant. **d.** Kaplan-Meier curves of OS per cohort, stratified with respect to the GTEx threshold (high score: activity above threshold, low score: activity below threshold). For mesothelioma, since no GTEx data could be matched with cancer data, the threshold is the median value. P-values were obtained with a log rank-test and corrected with Benjamini-Hochberg method to account for multiple-hypothesis testing.

After determining these indication-specific thresholds, we computed the proportion of patients with a “high” YAP1/TEAD activity (TEAD+ patients). As depicted in Figure 2a, this proportion ranged from 3% for prostate cancer (PRAD) to 98% for glioblastoma (GBM). These results are consistent with previous findings suggesting no evidence of a higher activity level in prostate cancer with respect to normal tissue^28^. Additionally, higher levels of *YAP1*/*TAZ* mRNA has been shown in glioma tissues^29^, in line with the high prevalence of TEAD+ patients in GBM observed in our study (Figure 2a).

Tumor types with a high proportion of TEAD+ patients included gynecologic cancers, such as uterine corpus endometrial carcinomas (UCEC, 53%), uterine carcinosarcomas (UCS, 91%), ovarian cancer (OV, 75%) and cervical squamous cell carcinoma (CESC, 66%). Other indications with intermediate levels of YAP1/TEAD activity encompassed various sarcomas (SARC, 64%), adrenocortical carcinoma (ACC, 53%) and head and neck squamous cell carcinoma (HNSC, 51%). At the other end of the spectrum, YAP1/TEAD activity level was very low in pancreatic adenocarcinoma (PAAD), pheochromocytoma and paraganglioma (PCPG), thyroid carcinoma (THCA) and kidney chromophobe cancer (KICH).

In a few indications, the signature exhibited a multimodal distribution, highlighting the presence of subgroups of interest. This was particularly clear in breast cancer (BRCA) where 78% (N=155) of patients were classified as TEAD+ in the subgroup of basal-type tumors, compared to the subgroups of HER2-enriched patients (20%, N=17), luminal B tumors (10%, N=24) and luminal tumors (only 0.3%, N=2, Figure 2b). In NSCLC, 17% (N=174) of TEAD+ tumors were observed overall. This proportion increased to 23% (N=124) in squamous cell carcinomas (LUSC) and to 52% (N=30) for patients belonging to the primitive subtype^30^. These results corroborate previous studies reporting higher YAP1/TEAD activation in squamous versus non squamous types^19^, as well as heterogeneous YAP1/TEAD activation across cancer subtypes, such as basal versus luminal A/B or HER2-enriched breast cancers^31^.

We also investigated the association between the TEAD-500 signature and patient survival. As expected, higher activity levels were observed in cancer types with poorer prognosis^19,20^. In particular, the fraction of TEAD+ patients correlated negatively with the 5-year survival rate computed for overall survival (OS) with R = -0.41 and p-value = 0.035 in all TCGA datasets tested (Figure 2c).

We then assessed the prognostic value of the TEAD-500 score in each cancer type by computing a univariate C-index^32^. P-values were corrected with Benjamini-Hochberg procedure to account for multiple testings (see Table S1 in supplementary data). The score was highly predictive of OS in MESO, with a univariate C-index of 0.72 (p = 1 × 10^−7^), higher scores being associated with a worse prognosis. Its value was significantly associated with poor prognosis in eleven other indications, the strongest associations being found in Adrenocortical carcinoma (ACC; C-index = 0.79, p = 3 × 10^−6^), KICH (C-index = 0.79; p = 7 × 10^−4^), Kidney Renal Papillary Cell Carcinoma (KIRP; C-index = 0.74, p = 3 × 10^−6^) and Lower Grade Glioma (LGG; C-index = 0.73, p = 5 × 10^−12^). Significant associations were also found in Liver Hepatocellular carcinoma (LIHC; C-index = 0.60, p = 7 × 10^−4^), SARC (C-index = 0.60, p = 0.011), Bladder Urothelial Carcinoma (BLCA; C = 0.59, p = 0.014), Lung Adenocarcinoma (LUAD; C-index = 0.56, p = 0.014), HNSC (C-index = 0.56, p = 0.005), Skin Cutaneous Melanoma (SKCM; C-index = 0.55, p = 0.017) and Kidney Renal Clear Cell Carcinoma (KIRC; C-index = 0.54, p = 0.004). Only in thymoma (THYM) were higher values associated with a better outcome (C-index = 0.67, p-value = 0.014). These results further validate the use of the TEAD-500 signature: for comparison, the signature defined in Wang et al.^19^ reached significance on eight indications, six of which are common to both signatures (LGG, MESO, KIRP, HNSC, BLCA and PAAD).

For each indication, we also examined the stratification of the population in two subgroups, based on the thresholds defined previously (with the exception of MESO where the median value was used) (Figure 2d). We computed the logrank p-values between patients with high and low YAP1/TEAD activity, and corrected the results with the Benjamini-Hochberg procedure. The stratification reached statistical significance in 6 indications out of 29: MESO (p = 2 × 10^−4^), LGG (p = 2 × 10^−5^), LIHC (p = 0.016), ACC (p = 0.004), SARC (0.010) and SKCM (p = 0.027) (Figure 2d).

Altogether, our data confirmed that YAP1/TEAD activity is both cancer type- and subtype-specific, and that abnormally high activity is correlated with poor prognosis in certain cancers.

### Deep learning can predict YAP1/TEAD activity from histology slides and can be used as a surrogate for pathway activity

To predict this activity signature on H&E-stained pathology slides, we developed a deep learning model called HE2TEAD, which relies on a weakly supervised learning framework. This approach is based on Chowder^33^, a model specifically designed to analyze Whole Slide Images (WSIs). HE2TEAD partitions the WSI into smaller patches – or tiles – measuring 112 × 112 µm each, that are first scored independently by the model. These tile-level scores are then aggregated to generate a prediction for the entire slide (architecture in the “Methods”). Training of the model was conducted on 18 separate cohorts from TCGA.

We first performed a five-fold cross-validation on each cohort. This involved randomly assigning patients to five different sets, where each set was alternatively used as the validation set, while the remaining four sets were used for training. The final results were expressed as a mean over all five runs (for the architecture, see “Methods”). Subsequently, the Pearson correlation (R) between the model prediction and the signature derived from bulk RNAseq (Figure 3a) was computed for each cohort.

**Figure 3.**
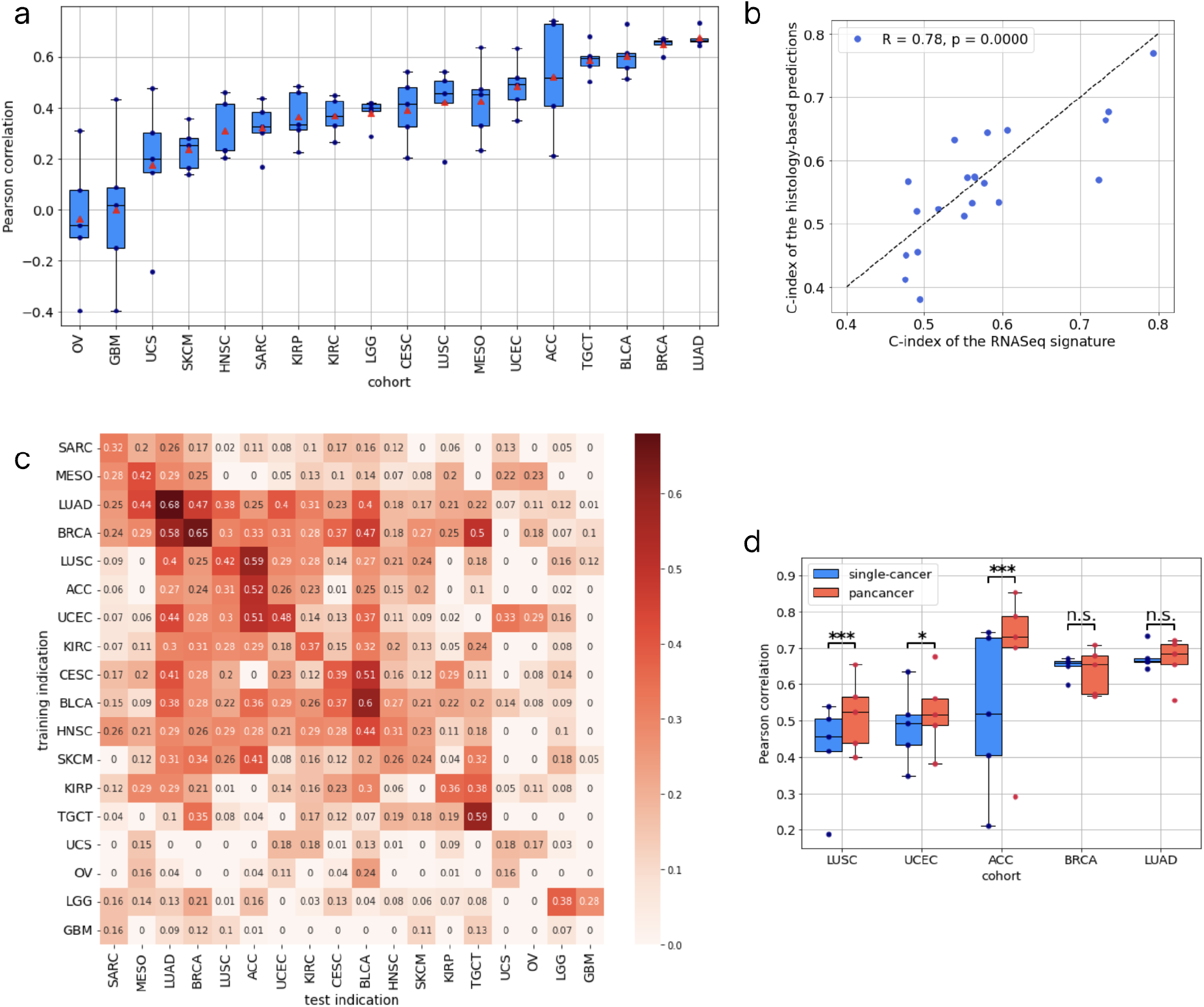
Deep learning can predict YAP1/TEAD activity from histology slides. **a.** Boxplot of the Pearson correlation values obtained in cross-validation on TCGA cohorts (box: interquartile range (IQR); horizontal line: median; whiskers: 1.5 times IQR, triangle: mean; circles: individual fold values). **b.** Correlation between the prognostic power of the RNAseq signature and that of the histology-based predictions. **c.** Table of the Pearson correlation obtained by applying a model trained on a given indication to any other indication. Diagonal values are the averages obtained in cross-validation. For readability, negative values were clipped to 0. The blue square indicates indications that were grouped for training the final NSCLC model **d.** Boxplot (defined as in panel **a**) of the Pearson correlation values obtained in cross-validation on LUSC, UCEC, ACC, BRCA and LUAD, with models trained individually on each cancer (blue) or with a single model trained simultaneously on the five cohorts (red). P-values were obtained from a Z-test, as described in the method section (* p-value < 0.05, ** p-value < 0.01, *** p-value < 0.001).

As presented in table 1, the models achieved high performances, with correlations exceeding 0.6, in BLCA (R = 0.603, std = 0.080), LUAD (R = 0.675, std = 0.050) and BRCA (R = 0.649, std = 0.047). The latter result is not surprising since the activation of the pathway strongly discriminates between two distinct subtypes of breast cancer, as mentioned previously (Figure 2b). The model was highly predictive of the BRCA subtypes, as indicated by the area under the curve (AUC) in distinguishing the basal subgroup from the other subtypes, namely luminal A and B and HER2-enriched subtypes (AUC = 0.88, std = 0.028). Despite the small size of the training dataset, HE2TEAD performed relatively well in Mesothelioma (R = 0.425, std = 0.153, with N = 95 samples). Conversely, the model’s performance was suboptimal in indications where the activation values were uniformly high (with most samples exceeding the threshold determined from healthy tissue), such as GBM, UCS and OV.

**Table 1:** Pearson correlation between the model’s predictions and the RNAseq-based signature.

|  | Single-cancer performance (Pearson R) |  |
| --- | --- | --- |
|  | average | standard deviation |
| ***LUAD (Lung Adenocarcinoma)*** | 0.675 | 0.050 |
| ***BRCA (Breast Cancer)*** | 0.649 | 0.047 |
| ***BLCA (Bladder Cancer)*** | 0.603 | 0.080 |
| ***TGCT (Testicular Germ Cell Tumors)*** | 0.588 | 0.064 |
| ***ACC (Adrenocortical Carcinoma)*** | 0.522 | 0.225 |
| ***UCEC (Uterine Corpus Endometrial Carcinoma)*** | 0.485 | 0.103 |
| ***MESO (Mesothelioma)*** | 0.425 | 0.153 |
| ***LUSC (Lung Squamous Cell Carcinoma)*** | 0.421 | 0.125 |
| ***CESC (Cervical Squamous Cell Carcinoma)*** | 0.392 | 0.133 |
| ***LGG (Lower Grade Glioma)*** | 0.380 | 0.053 |
| ***KIRC (Kidney Renal Clear Cell Carcinoma)*** | 0.368 | 0.074 |
| ***KIRP (Kidney Renal Papillary)*** | 0.363 | 0.107 |
| ***SARC (Sarcoma)*** | 0.323 | 0.102 |
| ***HNSC (Head and Neck Squamous Cell Carcinoma)*** | 0.309 | 0.119 |
| ***SKCM (Skin Cutaneous Melanoma)*** | 0.239 | 0.088 |
| ***UCS (Uterine Carcinosarcoma)*** | 0.175 | 0.265 |
| ***GBM (Glioblastoma)*** | -0.001 | 0.307 |
| ***OV (Ovarian Cancer)*** | -0.037 | 0.259 |

Since the TEAD-500 signature was associated with OS in several cancers, we checked whether the HE2TEAD prediction from the H&E slides retained this association with survival. We computed the concordance index on each cross-validation fold and averaged the results. In several indications such as ACC (C-index = 0.77, std = 0.12), KIRP (C-index = 0.68, std = 0.10), LGG (C-index = 0.66, std = 0.058), UCEC (C-index = 0.64, std = 0.047), BLCA (C-index = 0.56, std = 0.049) and LUAD (C-index = 0.57, std = 0.055), the prognostic power of the histology-based model was close to that of the TEAD-500 signature. Overall, the C-index obtained with the histology-based predictions was strongly correlated with the one obtained from the RNAseq signature (Pearson R = 0.78, p-value = 3 × 10^−5^).

Next, we searched for similarities in histological patterns between various cancer types. Our hypothesis was that if histological patterns associated with the TEAD+ phenotype were conserved in two indications, a model trained on one of those indications should transfer robustly to the other. To assess this, a model trained on a given cohort, was then applied to every other cancer type, each treated as an external validation dataset. We computed Pearson correlation and p-values, which were corrected for multiple hypothesis testing with the Benjamini-Hochberg method, thus obtaining a transfer table where rows represented the training indications and columns the test indications (Figure 3c).

The transfer matrix revealed significant off-diagonal values, highlighting similarities in the morphological patterns associated with YAP1/TEAD activity across several cancer indications. LUAD and BRCA appeared to be pivotal indications, since models trained on those cohorts transferred well to several other diseases: for instance, the BRCA-trained model achieved correlations as high as 0.58 on LUAD (p = 2 × 10^−47^), 0.47 on BLCA (p = 5 × 10^−25^) and 0.50 on TGCT (p = 5 × 10^−14^). BRCA and LUAD resided at the intersection of two groups of indications sharing a high similarity: one including MESO and SARC - in line with the known histological resemblance between mesothelioma and sarcoma^34^, and a second group with numerous cancers, including LUSC, ACC and UCEC, that could be extended to include all adenocarcinomas and squamous cell carcinomas. Among this extended group, BLCA and CESC formed a distinct subgroup (CESC to BLCA: R = 0.51, p = 10^−29^, BLCA to CESC: R = 0.37, p = 6 × 10^−10^). There was also a relatively good model transferability from UCEC to other gynecologic cancers (R = 0.33, p = 0.004 on UCS; R = 0.29, p = 0.014 on OV), and among brain cancers (LGG to GBM: R = 0.28, p = 3 × 10^−5^).

Several factors may explain those results. As an example, good transfer was observed among cancers affecting the same tissue or organ (LUSC and LUAD, UCEC and UCS, LGG and GBM) or with histological resemblance (MESO and SARC). Another potential contributing factor is the similarity in the underlying mechanisms of Hippo pathway inactivation. For instance, BLCA and CESC share a high frequency of mutations among the Hippo pathway genes (in particular *NF2*), as well as frequent amplification of *YAP1* and deletion of *LATS1*^19^. Based on this, we sought to investigate the efficacy of pancancer training of our models.

We retrained three models by grouping several TCGA cohorts together. Cross-validation was conducted on the resulting aggregated dataset. However, the performances were still evaluated separately on each cancer. We initially developed a model for the most closely related cancer types identified by the previous analysis: LUSC, LUAD, BRCA, UCEC and ACC, collectively referred to as the NSCLC model. Its average performance on these cohorts (R = 0.60) was significantly superior to that of single-cancer trained models (R = 0.55 , p = 2 × 10^5^; Figure 3d). To further optimize the performance on HNSC and MESO, we considered several combinations based on the transfer matrix and selected the one giving the best performance on those cohorts (see Methods for further details). We trained a model on HNSC, SKCM and BLCA (adding the two indications giving the best transfer performances on HNSC), in addition to the five previously mentioned indications. This model achieved a significant improvement (p = 0.001) with a correlation of 0.44 (std = 0.08) on TCGA-HNSC compared to 0.31 (std = 0.12) for the single-cancer trained model. Similarly, to enhance the performance on MESO, we trained a model jointly on LUAD and MESO. The resulting model exhibited a correlation of 0.48 (std = 0.16) on TCGA-MESO, higher, although not significantly so (p = 0.23), compared to the model solely trained on TCGA-MESO.

Finally, we applied these models on seven validation cohorts outside of TCGA, encompassing a total of 793 samples from patients with NSCLC (N= 419), HNSC (N=260), and MESO (N=114), as described in the Methods section (Figure 4).

**Figure 4.**
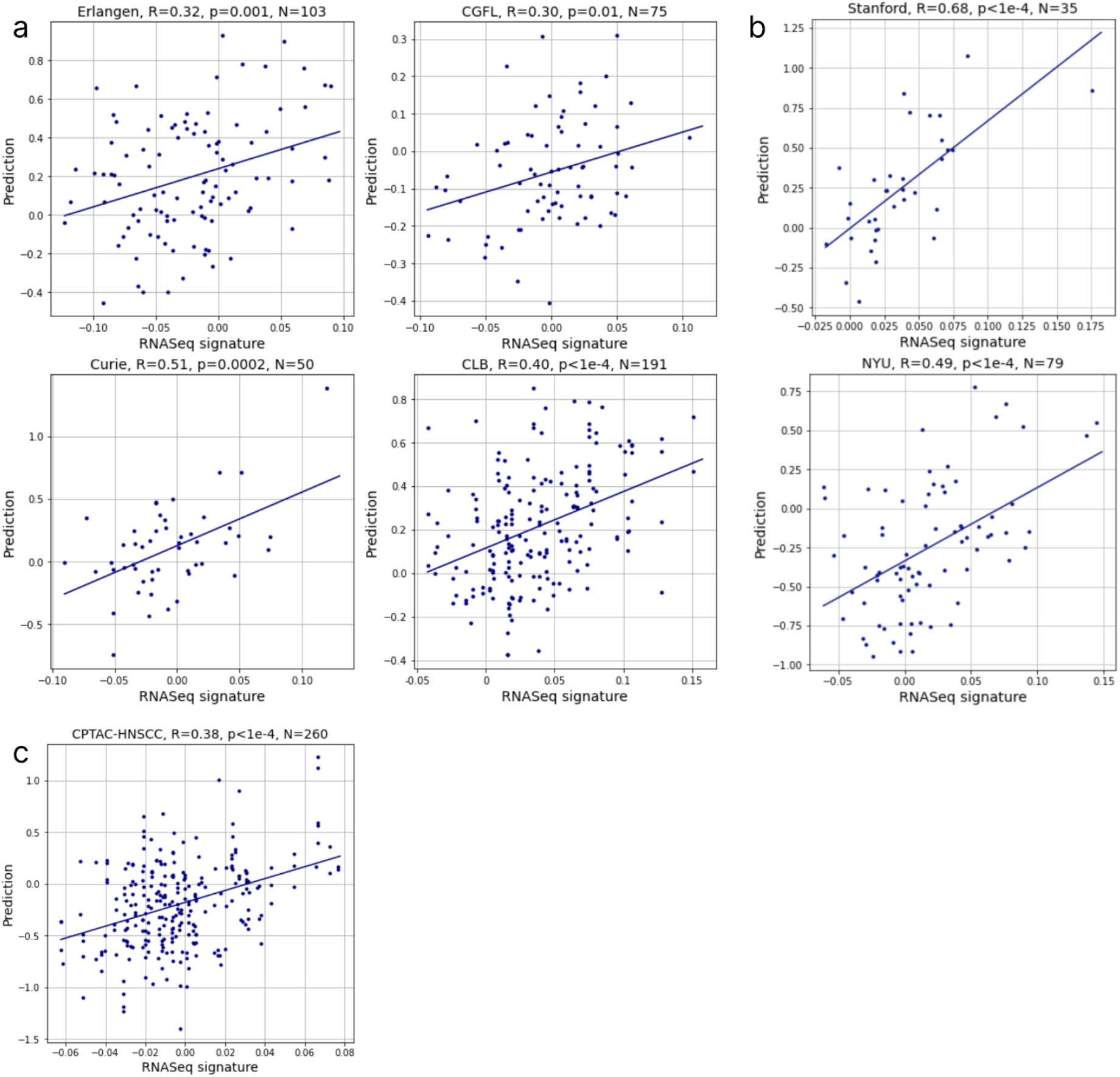
The HE2TEAD model transfers robustly to external validation cohorts. **a.** RNAseq signature values vs histology-based prediction on external NSCLC validation cohorts. Predictions were obtained from a model trained on advanced-stage patients (AJCC stage IV) from five TCGA cohorts: LUAD, LUSC, ACC, UCEC and BRCA. 2-sided p-values are reported. The scale is different on both axes because the model was trained to predict the normalized signature value (zero mean and unit standard deviation) **b.** Same as **a**, on external mesothelioma cohorts from Stanford and NYU. The model was trained on TCGA-MESO and TCGA-LUAD. **c.** Id., on the CPTAC-HNSCC cohort (head and neck squamous cell carcinoma). The model was trained on 7 TCGA cohorts: HNSC, SKCM, BLCA, ACC, BRCA, LUAD, LUSC, UCEC.

We validated the NSCLC model on four external cohorts, from Institut Curie (Curie), Centre Léon Bérard (CLB), Centre Georges-François Leclerc (CGFL) and Erlangen. Due to the limited number of patients with squamous cell carcinoma in each cohort, all histological subtypes were collectively analyzed. The model demonstrated predictive power on all external cohorts, with correlations ranging from 0.21 [0.03 - 0.39] (Erlangen, p-value = 0.032) to 0.49 [0.17 - 0.71] (Curie, p-value = 3 × 10^−4^). However, we observed a significant drop in performances compared to the cross-validation on TCGA (R = 0.58, std = 0.06 on TCGA-LUAD and TCGA-LUSC together) (Table 2).

**Table 2:** Pearson correlations and p-values of the trained models on the 4 available external cohorts of NSCLC. 95% Confidence Intervals (CI) were obtained by bootstrapping the cohort 10,000 times with replacement.

| Cohort | Model trained on TCGA (LUSC, LUAD, BRCA, UCEC and ACC) - all stages |  |  | Model trained on TCGA (LUSC, LUAD, BRCA, UCEC and ACC) - advanced stages |  |  |
| --- | --- | --- | --- | --- | --- | --- |
|  | Pearson R | 95% CI | p-value | Pearson R | 95% CI | p-value |
| Curie | 0.49 | 0.17 - 0.71 | $3 \times 10^{-4}$ | 0.51 | 0.17 - 0.72 | $2 \times 10^{-4}$ |
| CLB | 0.37 | 0.23 - 0.51 | $9 \times 10^{-8}$ | 0.40 | 0.27 - 0.52 | $1 \times 10^{-8}$ |
| CGFL | 0.29 | 0.11 - 0.45 | 0.011 | 0.30 | 0.11 - 0.46 | 0.010 |
| Erlangen | 0.21 | 0.03 - 0.39 | 0.032 | 0.32 | 0.14 - 0.47 | 0.001 |
| Average | 0.34 | 0.24 - 0.43 | $7 \times 10^{-11}$ | 0.38 | 0.27 - 0.47 | $2 \times 10^{-13}$ |

We hypothesized that this gap between TCGA cross-validation and external validation could be primarily attributed to differences in the patient populations. In particular, the external cohorts included large proportions of advanced-stage and metastatic cancers, whereas TCGA-NSCLC cohorts (LUAD and LUSC) were predominantly composed of primary tumors from patients with early-stage disease. To validate this hypothesis, we retrained a model on the same TCGA cohorts, exclusively selecting patients with an American Joint Commission on Cancer (AJCC) stage IV. This drastically reduced the size of the training dataset from 2,966 samples (1,034 NSCLC samples) to a mere 95 samples (33 NSCLC cases). The performances of this retrained model - referred to as TCGA-advanced - on external cohorts were improved on all cohorts, and particularly so on Erlangen (R = 0.32 [0.14 - 0.47], p-value = 0.001) (Table 2), although statistical significance was not reached (p = 0.08 for Erlangen, p = 0.27 across cohorts, Fisher test). It should be noted that the Curie and CLB cohort were stained with Haematoxylin, Eosin and Saffron (HES) while the training cohort (TCGA) only contained Haematoxylin and Eosin (H&E). The fact that good performances were achieved on those cohorts highlights the model’s robustness to staining variations.

The MESO model was validated on two external cohorts, from Stanford and NYU. In the Stanford cohort, the model was predictive of YAP1/TEAD activity with a correlation of 0.68 [0.51 - 0.81] (p-value = 7 × 10^−6^), while in the NYU cohort it achieved a correlation of 0.49 [0.29 - 0.64] (p-value = 5 × 10^−6^).

The HNSC model was validated on the CPTAC-HNSCC cohort^23^. Multiple RNAseq measurements were available for some patients, in which cases the replicate YAP1/TEAD activity scores were averaged across samples. The model achieved a correlation of 0.38 [0.27 - 0.48] (p-value = 2 × 10^−10^). These performances were comparable to those obtained in cross-validation.

Altogether, these results demonstrate that our models trained on TCGA are robust and can generalize to several external cohorts in different indications. This paves the way for potential clinical applications of our HE2TEAD models.

### YAP1/TEAD activity is associated with necrosis, poorly differentiated tumor and inflammation

Building upon the robustness of our models, we aimed to further characterize the histological patterns associated with YAP1/TEAD activity across cancer types. We selected the most predictive tiles of each model in several cohorts (TCGA-NSCLC, Erlangen-NSCLC and CLB-NSCLC, TCGA-HNSC and CPTAC-HNSCC, TCGA-MESO and Stanford-MESO). For each considered cohort, we extracted 100 tiles associated with a prediction of a high level of YAP1/TEAD activity, 100 associated with a low prediction and 100 random tiles for comparison. Those tiles were subsequently reviewed by pathologists to systematically assess the histological patterns associated with YAP1/TEAD activity (see Methods sections). For NSCLC, we selected the model trained on advanced-stage patients from TCGA, as it generalized better to external cohorts. Also, since late stage tumors have had more time to evolve either due to the expected evolutionary course of disease progression^35^ or to cell plasticity in response to treatment^36^, this model was expected to pick up more diverse features.

In all three TCGA cohorts (NSCLC-advanced, MESO and HNSC), the presence of necrosis was strongly associated with high levels of YAP1/TEAD activity. This association was also found in external cohorts, albeit to a lesser extent in the CLB and Erlangen NSCLC cohort (15% of tiles with positive predictivity in CLB, 22% in Erlangen, up to 69% in TCGA-HNSC). In Erlangen, the model also preferentially focused on poorly differentiated tumor areas (NOS tumor, p = 0.0001), which the pathologists could not explicitly classify as adenocarcinoma or squamous cell carcinoma by looking at the tile area only (Figure 5). Differences in pathological patterns associated with YAP1/TEAD activity between the NSCLC cohorts may be attributable to different sampling procedures and sample selection criteria. Indeed, TCGA only contains resection slides, whereas the CLB and Erlangen cohorts are mostly composed of biopsy specimens. The latter tend to contain less necrosis, as they are intended to primarily focus on viable tumor cell nests. We noticed a significant presence of hemorrhages in tiles from the Erlangen cohort (present in 27% of tiles randomly sampled from this cohort), which contrasts with observations from the other NSCLC datasets (only 2% and 1% of tiles randomly sampled from TCGA and CLB respectively). This difference is a likely consequence of the sampling technique, as most samples from Erlangen were obtained by endobronchial ultrasound scan and biopsy (EBUS), and might also explain why the models have lower performances on this particular cohort.

**Figure 5.**
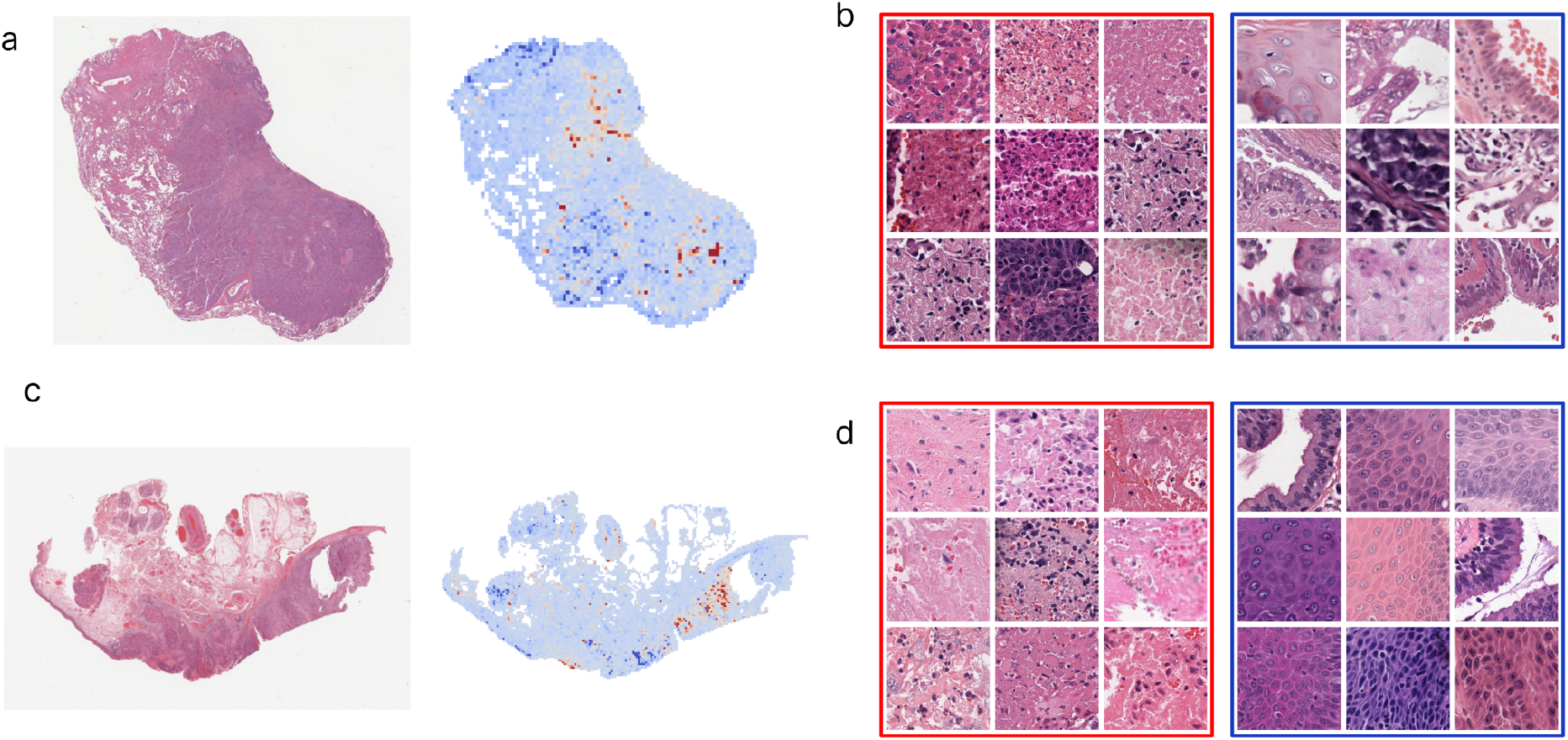
HE2TEAD identifies areas associated with a high YAP1/TEAD activity. **a.** Heatmap of the model on an NSCLC sample from TCGA. Red indicates regions associated with a high YAP1/TEAD activity value, while blue indicates areas associated with a low activity. **b.** Top positively predictive (red square) and negatively predictive tiles (blue square) from the TCGA-NSCLC cohort (advanced-stage patients). **c.** Heatmap of the model on an HNSC sample from TCGA. **d.** Top positively predictive and negatively predictive tiles from the TCGA-HNSC cohort.

**Figure 6.**
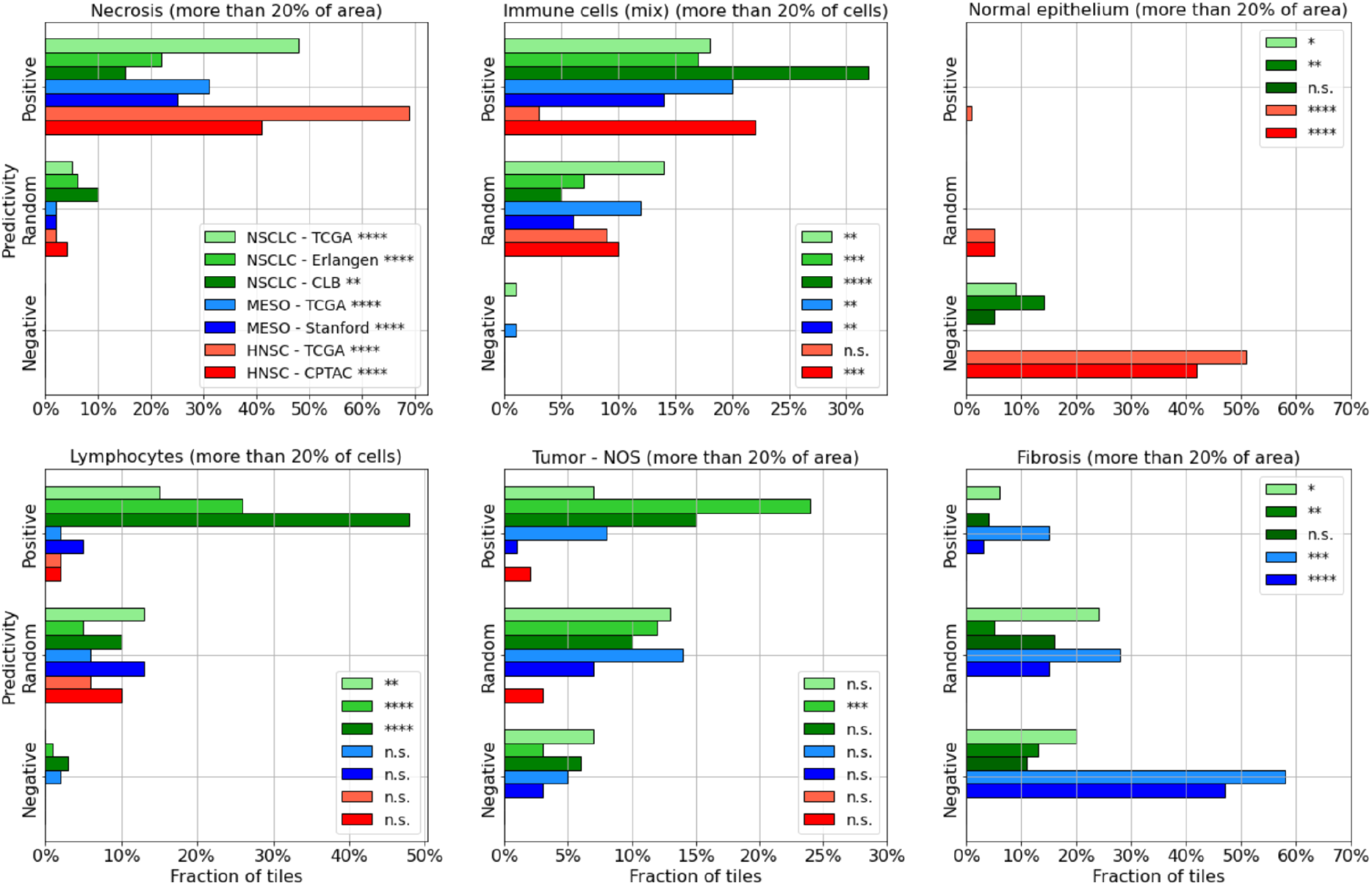
YAP1/TEAD activity is associated with necrosis and inflammation. Distribution of some of the histological patterns in positively and negatively predictive tiles from seven cohorts, as well as in randomly chosen tiles for comparison. P-values were obtained with a chi-square test of contingency between positive and negative tiles, and corrected for multiple-hypothesis testing with the Benjamini-Hochberg method. No tiles were annotated with normal epithelium in the mesothelioma cohorts, and similarly for fibrosis in the head and neck squamous cell carcinoma cohorts.

Furthermore, the presence of immune cells (exceeding 20% of the tile cell composition) was also associated with high YAP1/TEAD activity across most cohorts. Lymphocytes were found in predictive tiles from all NSCLC cohorts, with a very strong association in CLB (p = 2 × 10^−7^) and Erlangen (p = 1 × 10^−4^), while other immune cells (excluding lymphocytes) were predictive in all cohorts except TCGA-HNSC (Figure 5).

Altogether our findings are consistent with the known biological impact of the YAP1/TEAD signaling pathway in oncogenesis. Wang *et al.* recently established a strong association between hypoxia and the activation of the YAP/TAZ pathway. Their team also described immune cell abundance, such as dendritic cells in HNSC, MESO, and NSCLC, CD4 T cells and neutrophils in HNSC and NSCLC, macrophages in NSCLC, or B cells in MESO, to be associated with YAP/TAZ activity. Furthermore, several studies^37,38,39,40,41^ have pointed to the role of YAP1 and TAZ in promoting ferroptosis – an iron-dependent form of regulated necrosis – via the regulation of the *SKP2* gene^37^, one of the positive effectors of the TEAD-500 signature. Overall, this may be consistent with the preponderance of necrosis observed in all of the studied cohorts.

Presence of fibrosis or normal epithelium was predictive of low YAP1/TEAD activity. Up to 20% of the negatively predictive tiles presented fibrotic patterns in TCGA-NSCLC and this percentage reached almost 60% in TCGA-MESO. Additionally, normal epithelium was significantly correlated with low TEAD activity in 50% of the tiles in TCGA HNSC. Given that fibrotic tissues and normal epithelium are in essence non-tumoral, they are therefore expected to present lower YAP1/TEAD activity compared to the adjacent tumoral tissues.

All the above histological associations with YAP1/TEAD activity are consistent with well known roles of the Hippo pathway and its effectors in oncogenesis, namely the regulation of apoptosis and cell proliferation. Of particular pharmacological interest is the increased immune infiltration in tiles predicted to have a high YAP1/TEAD activity. This observation corroborates previous findings suggesting that an activated YAP/TAZ signaling promotes immune cell recruitment^42,19^.

## Discussion

Dysregulation of the Hippo pathway has been implicated in the oncogenesis of several cancers, as well as in resistance to therapies, which opens the way to translational research for the development of therapeutic agents targeting this pathway. Recently, numerous approaches have been developed across multiple modalities to disrupt the YAP1/TAZ–TEAD interaction^43^, resulting in several phase 1 clinical trials, primarily focusing on mesothelioma^44,45,46,47^.

However, extending this kind of therapeutic approach to other cancer indications requires tools to identify patient subpopulations that may benefit from the inhibition of YAP1/TAZ or TEAD. Here, we applied the RNAseq-based YAP1/TEAD activity signature defined in a recent study^10,20^ to every TCGA cancer type. Our analysis showed that the level of YAP1/TEAD activity is not only heterogeneous between different cancers, as previously documented in the literature, but can also display intra-indication variability, as evidenced for instance in breast cancer or in lung squamous cell carcinoma. In breast cancer in particular, the higher activity observed in the basal subtype is consistent with the role of hypoxia in this subtype^48^ and with the known association between hypoxia and Hippo pathway inactivation^49^. Moreover, our study confirms recent work indicating a correlation between Hippo pathway expression and hormone receptor status, notably estrogen receptor (ER) expression^31^.

The prognostic value of the signature was significant in 13 cancers out of the 33 TCGA indications, 12 of which showed a detrimental association with patient outcomes. This is also in agreement with previous findings^10,19,31^ and highlights the role of TEAD in cancer dynamics.

Although omics data have been used as the gold standard to identify abnormalities in the Hippo pathway, H&E-based prediction may offer broader applicability, given that histology slides are routinely available in clinical practice. In particular, this approach does not require any additional resource-consuming analyses. We designed deep learning models capable of predicting the signature directly from WSIs at the time of diagnosis for several cancer types. By training the models on several TCGA cohorts separately and testing on other cohorts, we highlighted groups of indications that shared common patterns. Our approach was extensively validated on multiple cohorts of mesothelioma, NSCLC and HNSCC outside of TCGA, demonstrating its robustness with respect to various acquisition protocols and batch effects.

Our image-based models further revealed morphological changes associated with abnormal YAP1/TEAD activity. In particular, necrosis emerged as an important cross-disease marker, at least in the 3 cancer types considered in the present work. Further analyzes would be needed to reach a finer comprehension of disease-specific and subgroup-specific biomarkers. Overall, our results present high clinical relevance, as these interpretable biomarkers associated with YAP1/TEAD activity could help clinicians by facilitating patient diagnosis and by defining treatment inclusion criteria. Moreover, they can be more easily presented and explained to patients.

These results open the way to potential applications in clinical practice. For instance, image-based biomarkers may facilitate the recruitment of patients for clinical trials involving treatments based on TEAD inhibitors. By leveraging the H&E slides routinely generated during the patient treatment pathway, this approach is much more cost-effective than relying on the generation of RNA-based biomarkers. Moreover, it could accelerate diagnosis and treatment decision-making by clinicians as these analyses can be performed using the H&E slides obtained at the time of diagnosis. However, such applications would require additional developments that are beyond the scope of the present work.

## Acknowledgements

We thank The Cancer Genome Atlas (TCGA) program for providing free access to histology slide images and RNA-Seq data. We thank Donald Jackson, Jean-Philippe Vert and Jonas Béal for insightful discussions.

## Author contributions

B.S., V.C., O.D.D. and J.E.L.D. designed the experiments; P.M. performed the double-staining experiments; B.S., V.C., O.D.D. and J.E.L.D. performed the numerical experiments and wrote the code to achieve the different tasks; B.S., V.C., O.D.D., J.E.L.D, A.H., R.B., C.M., A.R., E.R., I.V., L.D., M.Cl., K.V.L., E.P. and M.Ce. analyzed the results.

B.S., V.C., O.D.D., J.E.L.D., A.H., R.B. and E.P. wrote the manuscript with the assistance and feedback of all the other co-authors.

## Declaration of interests

The authors declare the following competing interests:

### Employment

B.S., V.C., O.D.D., J.E.L.D, A.H., R.B., C.M., A.R., E.D., K.V.L. and E.P. are employed by

Owkin, Inc.; E.R., I.V., L.D., E.d.R., F.N., E.S., V.F., M.Cl. and M.Ce. are employed by Sanofi

### Advisory

M.E.: Personal fees, travel costs and speaker’s honoraria from MSD, AstraZeneca, Janssen-Cilag, Cepheid, Roche, Astellas, Diaceutics, Owkin, BMS; research funding from AstraZeneca, Janssen-Cilag, STRATIFYER, Cepheid, Roche, Gilead, Owkin; advisory roles for Diaceutics, MSD, AstraZeneca, Janssen-Cilag, GenomicHealth, Owkin, BMS.

A.S.: Honoraria for advisory boards: Amgen, Astrazeneca, Boehringer Ingelheim, Ipsen, Janssen, Lilly, MSD, Pfizer, Roche, Takeda. Consulting: Astrazeneca, BMS, Daiichi Sankyo Inc, Janssen, Roche. Symposiums: Amgen, Astrazeneca, BMS, Janssen, Pfizer, Sanofi, Takeda. Congress: Janssen, Pfizer, Takeda.

S.L.: Honoraria for advisory boards: Janssen, MSD, Sanofi, Abbvie. Symposiums: MSD, Janssen. N. G.: Honoraria and/or consulting fees from AbbVie, Amgen, AstraZeneca, BMS, Boehringer-Ingelheim, Daachi, Janssen, Lilly, Mirati, MSD, Novartis, Novartis, Pfizer, Roche, Sanofi, Takeda, and received grants from MSD and AstraZeneca paid to institution outside of the submitted work.

### Patents

E.S. holds a patent for the TEAD500 signature (WO2023180385A1).

## Data availability

The TCGA dataset is publicly available via the TCGA portal (https://portal.gdc.cancer.gov). The CPTAC dataset is publicly available via the TCIA portal (https://www.cancerimagingarchive.net/collection/cptac-hnscc/). Restrictions apply to the availability of the other datasets, which were used with permission for the current study, and so are not publicly available. Data can, however, be made available upon reasonable request from Erlangen University Hospital, Centre Léon Bérard (CLB), Centre Georges-François Leclerc (CGFL), New York University (NYU), Stanford University and Institut Curie. Specific conditions and restrictions of access to the datasets are to be discussed directly with the main investigators in each center: M.E. for Erlangen, A.S. for CLB, FGF. for CGFL, H.P. for NYU, O.G. for Stanford and N.G. for Institut Curie.

## Methods

### Data

This study was based on publicly available data from TCGA (https://portal.gdc.cancer.gov/). We selected samples from primary tumors only, for which both RNA-Seq and WSI data were available. Transcriptomic data (FPKM-UQ) were extracted from frozen tissues, and the slides analyzed were digitized H&E-stained formalin-fixed, paraffin-embedded (FFPE) histology slides, referred to here as whole-slide images (WSIs).

RNA-Seq samples from healthy tissue were obtained from the public database GTEx (https://gtexportal.org/home/datasets).

For external validation of our models, we had access to 4 NSCLC cohorts, 2 mesothelioma cohorts and one cohort of head and neck squamous cell carcinoma. The HNSCC validation cohort is publicly available from the CPTAC database (https://proteomics.cancer.gov/programs/cptac). All validation cohorts are described in table 3.

**Table 3:**
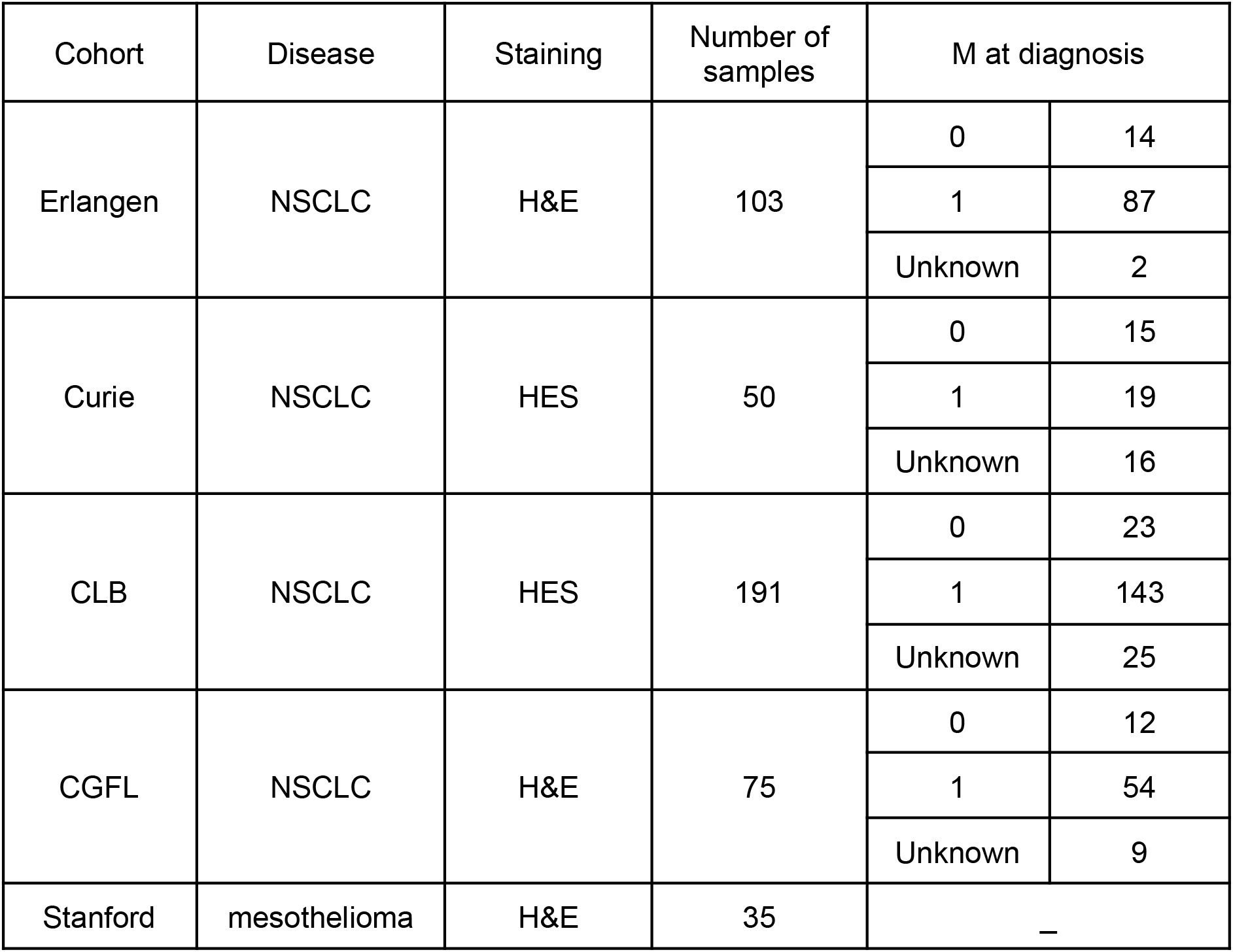
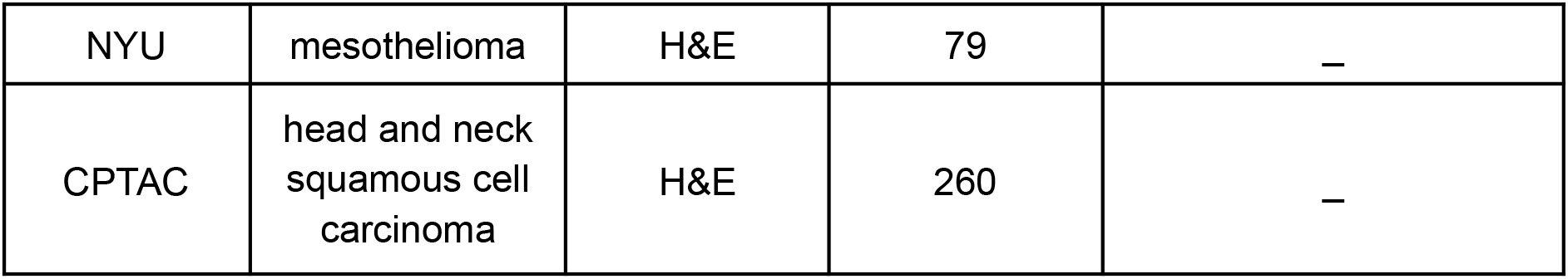
Description of the external cohorts used for validation of our models. For cohorts where the information was available, we report the metastatic status (M at diagnosis).

| Cohort | Disease | Staining | Number of samples | M at diagnosis |  |
| --- | --- | --- | --- | --- | --- |
| Erlangen | NSCLC | H&E | 103 | 0 | 14 |
|  |  |  |  | 1 | 87 |
|  |  |  |  | Unknown | 2 |
| Curie | NSCLC | HES | 50 | 0 | 15 |
|  |  |  |  | 1 | 19 |
|  |  |  |  | Unknown | 16 |
| CLB | NSCLC | HES | 191 | 0 | 23 |
|  |  |  |  | 1 | 143 |
|  |  |  |  | Unknown | 25 |
| CGFL | NSCLC | H&E | 75 | 0 | 12 |
|  |  |  |  | 1 | 54 |
|  |  |  |  | Unknown | 9 |
| Stanford | mesothelioma | H&E | 35 | — |  |

|  |  |  |  |  |
| --- | --- | --- | --- | --- |
| NYU | mesothelioma | H&E | 79 | – |
| CPTAC | head and neck squamous cell carcinoma | H&E | 260 | – |

### YAP1/TEAD activity score

There is no unique way of defining the activity score of a pathway from the transcriptome. Wang *et al.* compared 9 different measures of the YAP1/TAZ activity, derived either from the transcriptome or from protein assays, on the basis of their prognostic power of patient survival across cancer types. The authors reasoned that dysregulation of the Hippo pathway should be detrimental to the patient’s survival, and chose the score that was most consistently and extensively correlated with patient survival, a manually curated signature of 22 downstream genes of YAP1/TAZ significantly associated with survival in 8 TCGA indications.

In our study, we used Calvet et al’s YAP1-TEAD activity score, TEAD-500 signature, based on a list of 500 downstream effector genes derived from genetic perturbation experiments data instead of relying on prior knowledge. Even though it was not optimized for this task, it was shown to be even more associated with survival (13 indications including 12 where the signature is associated with poor outcomes).

To compute this score, the expression levels of the effector genes were ranked, and the difference of effector ranks (deR) score was defined as Rp – Rn, where Rp is the mean fractional rank of the positive effectors, and Rn, that of the negative ones.

### Preprocessing of RNAseq data

We used a reproducible in-house preprocessing pipeline to produce TPM count matrices from raw FASTQ files, as outlined below.

The preprocessing pipeline was developed as a Snakemake workflow^50^. First, FASTQ file trimming (if required) is performed using Cutadapt (v4.1). The reads were aligned on the GRCh38 reference genome assembly (downloaded from the “Broad References” dataset) using the STAR aligner (v2.7.9a). The read quantification was also performed by STAR, which is equivalent to HTSeq-count. Raw read counts were then normalized to TPM counts using known exonic gene sizes. TPM counts can be computed more precisely taking into account the proportion of different exons for each gene, but comparison shows almost no difference regarding the Calvet score (>99% correlation, same range of values).

Quality control (QC) was performed before (FASTQC) and after (RSeQC v4.0) alignment, and outliers were identified by performing PCA on raw counts, TPM, and DESeq2-normalized counts. Samples with less than 5 million reads and outliers were removed before proceeding to TEAD-500’s score computation using the TPM counts.

Mesothelioma samples are generally expected to be of lower quality compared to other tumoral samples due to the difficulty of the biopsy and the (usually) longer conservation of samples. Since the Calvet et al’s formula is quite robust to low quality samples, we used less stringent QC filters on MESO samples, with the following results:

- NYU: out of 93 total samples we removed 11 samples with less than 1M reads total.

No outliers were detected.

- Stanford: out of 42 samples we removed 3 samples with less than 5M reads total. No outliers were detected.

For public datasets (TCGA, GTEx and CPTAC), we used the raw counts directly rather than the FASTQ files and the preprocessing step was limited to the generation of TPMs using known exonic gene lengths, in order to be consistent with the in-house preprocessing pipeline.

### Preprocessing of histology data

Since whole-slide images (WSIs) can have very large dimensions (up to 100,000 x 100,000 pixels), they need to be processed with specialized techniques. We applied three preprocessing steps to reduce dimensionality and remove potential artifacts.

We first detected the part of the image that contains matter. This segmentation was performed using a UNet neural network^51^. The UNet’s encoder is a VGG11, pretrained on the ImageNet dataset, while the decoder is trained from scratch. The architecture was fine-tuned on an internal dataset of 470 manually segmented slides, with various stainings and acquisition conditions. All pixels were separated between two classes: tissue, and background or artifacts (such as pen marks for instance).

We divided the regions of the slides for which matter was detected into smaller images, called ‘tiles’, of fixed size (224 × 224 pixels at a resolution of 0.5 micron per pixel). At least 60% of the area of a tile had to be detected as foreground by the UNet to be considered as containing tissue. The number of tiles n_tiles_ depended on the total area of tissue detected and varied from a few thousands to almost a hundred thousand tiles for each slide.

Features were extracted from each tile image using a pretrained convolutional neural network (the “extractor”), that encoded each tile image into a numerical feature vector. Here, we used the self-supervised algorithm MoCo v2^52,53^ to train a feature extractor without supervision on unlabeled histology images. The training procedure generated multiple views of a specific tile and tried to extract mutual information between those views. It also generated an embedding from negative examples (other tiles in the dataset) as different as possible from the embedding of the positive example. To achieve this dual task, the model needed to learn the semantic meaning inside each of these views, thus creating a feature extractor that was tailored for histology. The extractor we used was trained on the TCGA-COAD cohort, as recent works^54,55^ have shown that a network pretrained on this cohort had good generalization capabilities. At the end of this step, each slide was represented by a matrix of n_tiles_ × 2,048 features.

### WSI models

WSI models were based on the Chowder architecture^33^. The model was applied directly to the feature vectors extracted at the previous stage. It consisted essentially of two steps:

- Top and negative instances: We used a convolutional one-dimensional (1D) layer to compute a score for each tile. This convolutional layer performed a weighted sum between all 2,048 features of the tile to obtain this score (weights of this sum were learned by the model). We then picked the 25 highest and the 25 lowest scores and used them as input for our last step. This architecture specifies which tiles are used to make the predictions and, therefore, how our algorithm predicts the result. In particular, this method allowed us to interpret high-score tiles as areas indicative of a positive prediction (i.e., in our case high TEAD activity) and low-score tiles as areas indicative of a negative prediction.

- Multi-layer perceptron (MLP) classifier: This step consisted of an MLP with two fully connected layers of 200 and 100 neurons with sigmoid activation. This is the core of the predictive algorithm that transformed the scores from the tiles into a prediction.

Compared with the original paper, we considered a simple modification of the model architecture: instead of computing only one score per tile, one can use a 1D convolution with several output channels. This allowed the model to capture richer patterns, as each score could encode a specific histological structure associated with the activation.

As noted in the original work, this architecture is highly sensitive to model initialization, and ensembling N copies of the same architecture with different seeds increased the robustness and the performances of the model.

For single-cancer models, we did not run an extensive optimisation of hyperparameters and used models with one convolution channel, and N = 50. The MESO model, trained on MESO and LUAD, has the same architecture. The NSCLC and HNSC models trained on several cancer types had 5 convolution channels, and N = 50.

For HNSC and MESO, we considered several possible groups of indications for training the models, and selected the ones that gave the highest average correlation in cross-validation on the cohort of interest.

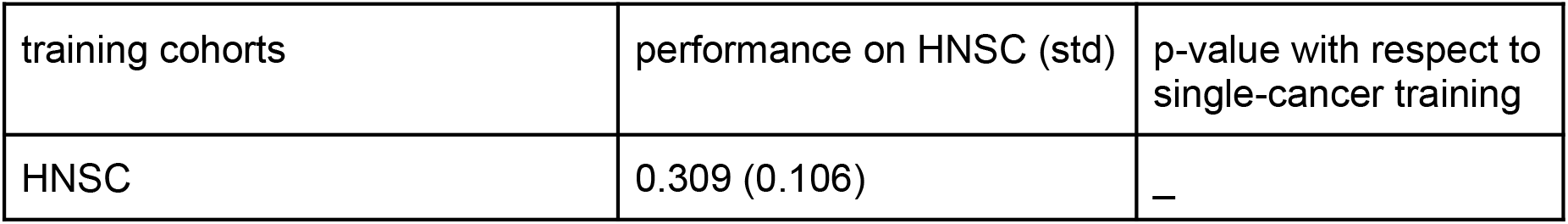
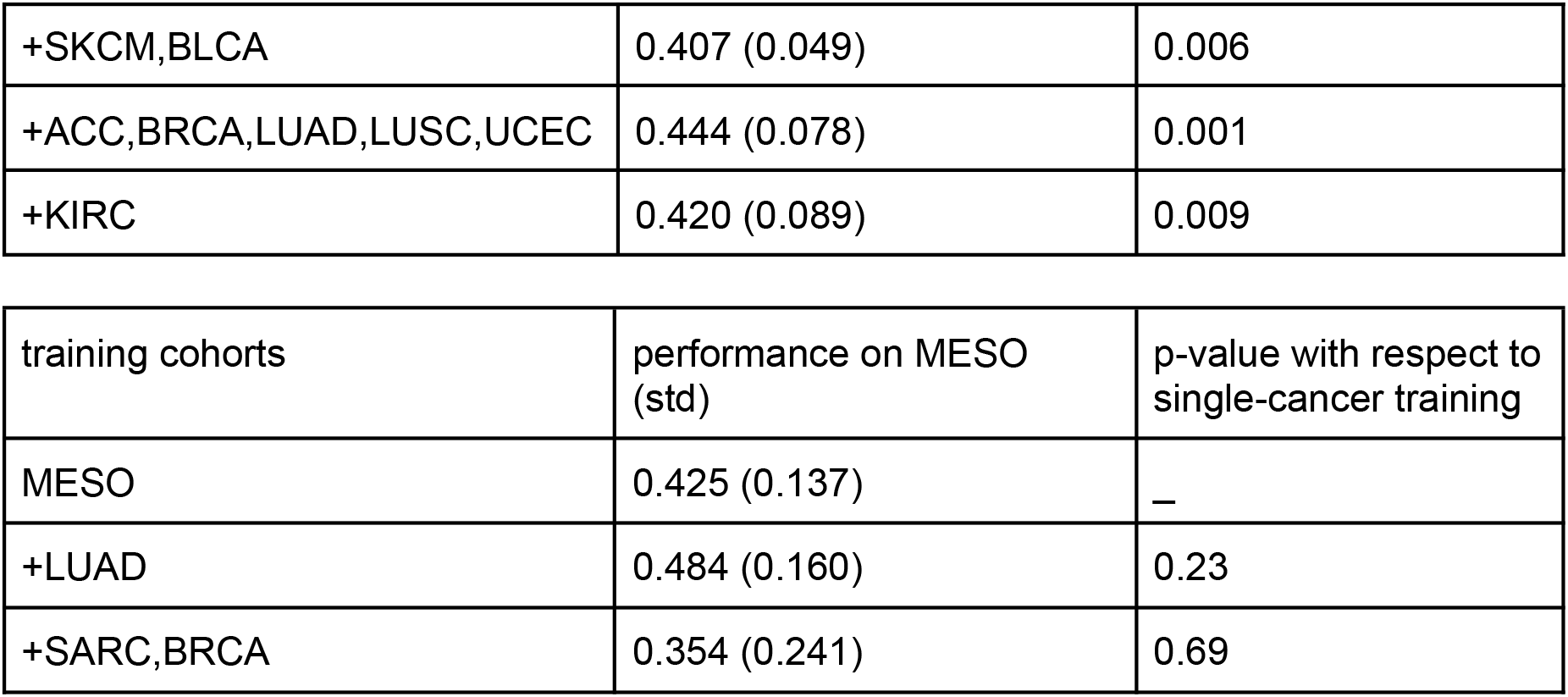

| training cohorts | performance on HNSC (std) | p-value with respect to single-cancer training |
| --- | --- | --- |
| HNSC | 0.309 (0.106) | — |
| +SKCM,BLCA | 0.407 (0.049) | 0.006 |
| +ACC,BRCA,LUAD,LUSC,UCEC | 0.444 (0.078) | 0.001 |
| +KIRC | 0.420 (0.089) | 0.009 |

| training cohorts | performance on MESO (std) | p-value with respect to single-cancer training |
| --- | --- | --- |
| MESO | 0.425 (0.137) | — |
| +LUAD | 0.484 (0.160) | 0.23 |
| +SARC,BRCA | 0.354 (0.241) | 0.69 |

### Model validation

For validation on external cohorts, all models trained in cross-validations were applied, and their predictions were averaged for each sample.

### Interpretability

As mentioned in the original paper, Chowder is interpretable by design due to the following two points.

- Only a small portion of the image is actually used by the model to make a prediction. Hence, once a model is trained, the predictive tiles can be readily accessed for analysis.

- Moreover, every tile obtains a score that can readily be interpreted as the local prediction of the model.

This is true for the baseline architecture, with a unidimensional score and no ensembling. However, since we used a multi-scoring approach and ensembling to increase the performances, interpretability becomes more challenging.

Instead, we adapted Shapley values^56^ to the specific case of histology models. In their original formulation, Shapley values addressed the distribution of gains to players in cooperative game theory. They have been applied in machine learning as a tool to interpret black box models. The application to our histology pipeline was quite straightforward. Slides were represented here as collections, or bags, of tiles, which is equivalent to a coalition of players in game theory, and we wanted to assess the contribution of each tile to the model’s prediction (equivalent to the gain). To do so, we considered every possible way of subsampling the tiles from a WSI. For each subsampling, we obtained a bag of n tiles (n varying from 1 to the total number of tiles in the WSI), and we computed the prediction of the model for this bag.

Then, the contribution of a given tile t_i_ was given by the difference between the average of the predictions obtained for bags containing t_i_ and the average of the predictions obtained for bags that do not contain t_i_. This exact formula was intractable in practice, since a WSI can contain up to 50,000 tiles, which would give 2^50,000^ possible bags. A way to overcome this was to randomly sample enough bags to obtain a reliable estimation. In practice, we sampled a maximum of 10,000 random bags and interrupted the computation in case of early convergence.

We checked that, when applied to the baseline architecture, Shapley values and Chowder scores gave consistent estimates of tile importance (in particular the most predictive tiles remained the same). Shapley values were a monotonously increasing function of Chowder scores, and tiles with intermediate scores (meaning they were not selected for the final prediction) had close to 0 contribution.

### Pathologist review

For the blind review by a pathologist, we selected 100 tiles which were the most associated with a high TEAD activity from the entire cohort (with a maximum of 3 tiles coming from the same slide). Similarly, 100 tiles associated with a low activation were selected. We also selected 100 random tiles in the whole cohort, again with a maximum of 3 tiles coming from the same slide. Those tiles were mixed and reviewed blindly by a pathologist, who also had access to a larger context: each tile had dimensions 112 x 112 µm, and we provided a context of size 1008 x 1008 µm. The pathologist had to evaluate each tile according to a predefined grid.

In order to standardize practices among pathologists, it was specified that tissue-level patterns (tumor, stroma, fibrosis…) had to cover at least 20% on a tile to be considered present. For cell-level patterns, the decision threshold was 20% of the cells in the tile, except for normal mesothelium and red blood cells where it was set at 20 cells in total.

| Cancer | Histological pattern |
| --- | --- |
| Mesothelioma | Tumor - epithelioid (more than 20% of area) |
|  | Tumor - sarcomatoid (more than 20% of area) |
|  | Normal mesothelium (more than 20 cells) |
| NSCLC | Tumor - adenocarcinoma (more than 20% of area) |
| NSCLC and HNSC | Tumor - squamous cell carcinoma (more than 20% of area) |
| All cancers | Tumor - NOS (more than 20% of area) |
|  | Stroma - collagen-rich (more than 20% of area) |
|  | Stroma - immune-rich (more than 20% of area) |
|  | Stroma - NOS (more than 20% of area) |
|  | Necrosis (more than 20% of area) |
|  | Hemorrhage (more than 20% of area) |
|  | Immune cells (mix) (more than 20% of cells) |
|  | Lymphocytes (more than 20% of cells) |
|  | Red blood cells (more than 20 cells) |
|  | Cartilage (more than 20% of area) |
|  | Fat tissue (more than 20% of area) |
|  | Muscle tissue (more than 20% of area) |
|  | Fibrosis (more than 20% of area) |
|  | Normal epithelium (more than 20% of area) |
|  | Artefact |

### Statistics and p-values

For cross-validation, average and standard deviation of the cross-validation metrics (Pearson correlation) are reported. For external validation, 95% confidence intervals were computed by bootstrapping, ie. drawing 10,000with-replacement samples from the dataset, computing the validation metrics for each bootstrap, and extracting the 2.5th and 97.5th percentile of the distribution thus obtained. We also reported 2-sided p-values, as implemented in methods stats.pearsonr of the python library scipy^57^. When comparing the performances of two models, we applied a Z-test on the average difference between the bootstrapped estimates. To aggregate comparison p-values on multiple independent test sets, we used Fisher’s method, as implemented in the python library scipy (scipy.stats.combine_pvalues).

To compare the proportion of tiles containing each histological pattern, we performed a chi-square test of proportions between positively predictive and negatively predictive tiles.

